# Viral susceptibility across host species is largely independent of dietary protein to carbohydrate ratios

**DOI:** 10.1101/2020.10.21.348987

**Authors:** Katherine E Roberts, Ben Longdon

## Abstract

The likelihood of a successful host shift of a parasite to a novel host species can be influenced by environmental factors that can act on both the host and parasite. Changes in nutritional resource availability have been shown to alter pathogen susceptibility and the outcome of infection in a range of systems. Here we examined how dietary protein to carbohydrate altered susceptibility in a large cross infection experiment. We infected 27 species of Drosophilidae with an RNA virus on three food types of differing protein to carbohydrate ratios. We then measured how viral load and mortality across species was affected by changes in diet. We found that changes in the protein:carbohydrate in the diet did not alter the outcomes of infection, with strong positive inter-species correlations in both viral load and mortality across diets, suggesting no species by diet interaction. Mortality and viral load were strongly positively correlated, and this association was consistent across diets. This suggests changes in diet may give consistent outcomes across host species, and may not be universally important in determining host susceptibility to pathogens.

**Twitter summary:** No role of host diet in susceptibility to a novel viral pathogen across host species

**Impact Statement:** A successful host shift of a parasite from one susceptible species to a novel host can be influenced by many ecological factors. Changes in host diet can alter the immune response and outcomes of host–parasite interactions and could potentially alter the outcome of a virus host shift. To investigate, we infected 27 species of Drosophilidae with an RNA virus (DCV) across three food types with differing protein to carbohydrate ratios. We then looked at pathogen loads and survival of infected hosts compared to uninfected controls. Changes in the ratio of protein to carbohydrate did not alter susceptibility to DCV across host species.

## Introduction

A key driver of pathogen host shifts – where a pathogen jumps from one host species to another – is environmental change (Hoberg & Brooks, 2015; Carlson *et al*., 2020). For a host shift to successfully occur a novel host must first be exposed to a parasite, which must then be able to replicate and successfully infect the host, before sufficient onward transmission (Woolhouse *et al*., 2005). Ecological factors can therefore influence the likelihood of host shifts by altering species distributions and abundances making encounters more likely, or by acting as stressors that alter physiological factors including immunity or virulence. The main ecological factor studied has been temperature, which can have asymmetrical impacts on hosts and parasites and potentially alter the likelihood of host shifts (Brooks & Hoberg, 2007; Hoberg & Brooks, 2015; Kirk *et al*., 2018; Roberts *et al*., 2018). The role of other ecological traits such as resource availability, humidity, population density and geographical range, or within host ecological traits such as metabolic rate, have been less well studied in explaining the outcomes of host shifts, despite an increasing understanding of the role such factors play in effecting the outcomes of host parasite interactions (Blanford & Thomas, 1999; Harvell *et al*., 2002; Ponton *et al*., 2013; Hayman *et al*., 2016; Cumnock *et al*., 2018).

Nutrition can shape the outcome of host-parasite interactions through its ability to moderate both parasite virulence and host resistance (Ponton *et al*., 2011, 2013; Pike *et al*., 2019). The nutritional resources of a host can impact its ability to resist infection as immune responses are thought to be costly to both maintain and activate (Kraaijeveld & Godfray, 1997; Lochmiller & Deerenberg, 2000; McKean *et al*., 2008; Cotter *et al*., 2011; Kutzer & Armitage, 2016; Knutie *et al*., 2017). Nutrition is known to have long term consequences, with developmental nutritional status being shown to have latent or even trans-generational effects on immune responses (in *Drosophila*: Fellous & Lazzaro, 2010; Savola *et al*., 2020b and reviewed in Grueber *et al*., 2018). Hosts can also show behavioural modifications in feeding upon infection; parasite-induced anorexia is thought to be an adaptive host response (Ayres & Schneider, 2009; Rao *et al*., 2017). In some cases hosts actively increase the consumption of certain nutrients in their diet for example, the African armyworm – *S. exempta* upon infection with a baculovirus displays macronutrient self-medication (Povey *et al*., 2009). Nutrition may constrain the amount of investment that a host can allocate to an energetically demanding immune response (Kraaijeveld & Godfray, 1997; Lochmiller & Deerenberg, 2000; Cotter *et al*., 2011; Knutie *et al*., 2017), and coping with costs associated with a parasite burden if infection does become established (Sheldon & Verhulst, 1996). A suboptimal nutritional status may lead a host to be unable to suppress or tolerate a parasite challenge they may otherwise have been able to resist; or have reduced fitness from a trade off in resources with life history traits (Kraaijeveld & Godfray, 1997; Lochmiller & Deerenberg, 2000; Cotter *et al*., 2011; Knutie *et al*., 2017).

From a parasite perspective infecting a host of suboptimal nutritional status may mean they encounter a weaker immune response and therefore infection and establishment is easier (Siva-Jothy & Thompson, 2002). However, once established the parasite may encounter its own resource limitations due to competition with an already depleted host, causing suboptimal growth and potentially affecting onward transmission. Therefore, predicting the outcome of the interaction between nutrition, host immunity and subsequent resistance is complex as the effects on the two parties may be divergent (Bedhomme *et al*., 2004).

Multiple life history traits are moderated by resource availability, with condition dependency across reproductive traits, aging and lifespan (Lee *et al*., 2008; Maklakov *et al*., 2009; Camus *et al*., 2017; Henry & Colinet, 2018; Henry *et al*., 2020). Laboratory experiments on dietary restriction, where individuals experienced a reduction in nutrition without inducing malnutrition (differentiated from Calorie Restriction) have been found to extend life span in a range of organisms (Weindruch & Walford, 1982; Klass, 1983; Anderson *et al*., 2003; Nakagawa *et al*., 2012). The effects of dietary restriction appear to be explained by resource-mediated trade-offs between longevity and reproductive effort (but see review by (Moatt *et al*., 2020)). Geometric frameworks – the use of artificial diets with known compositions of specific nutrients that develop an understanding of dimensional nutrient space – have been used to examine the consequences of different ratios of macronutrients across a range of organisms(Simpson & Raubenheimer, 1995, 2011; Raubenheimer & Simpson, 1999). In *Drosophila* different life-history traits were optimized at different protein-carbohydrate intakes (Lee *et al*., 2008; Skorupa *et al*., 2008; Fanson *et al*., 2009; Jensen *et al*., 2015). Across multiple species, low protein to carbohydrate ratios reduce reproductive rates but maximise lifespan (Nakagawa *et al*., 2012; Le Couteur *et al*., 2016). However, individuals with diets higher in protein and lower in carbohydrates have greater reproductive rates but shorter life spans. When given a choice of diet, individuals have been shown to optimise reproduction over lifespan (Bunning *et al*., 2016). Host dietary frameworks have been used to examine effects on bacterial pathogens (Povey *et al*., 2009; Cotter *et al*., 2019; Savola *et al*., 2020b; Wilson *et al*., 2020), viral pathogens (Lee *et al*., 2006; Povey *et al*., 2014), and individual aspects of immunity and gene expression (Cotter *et al*., 2011, 2019; Keaton Wilson *et al*., 2019). In particular, studies of viral infection in insects have found that high dietary protein leads to increased resistance (Lee *et al*., 2006) indicating there may be higher protein costs of resistance.

To investigate the effect that host diet has on the susceptibility of different host species we infected 27 species of Drosophilidae, with Drosophila C Virus (DCV) fed on three diets with varying ratios of protein to carbohydrates but comparable calorie content. We then measured the change in viral load and host mortality across these different diets. DCV is a positive sense RNA virus in the family *Dicistroviridae*. DCV was isolated from *D. melanogaster* although has also been detected in the closely related *D. simulans* (Christian, 1987), and in the wild it is thought to be transmitted faecal-orally (Jousset *et al*., 1972). Infection of DCV by inoculation is highly pathogenic in adult flies causing increased mortality rates, metabolic and behavioural changes and nutritional stress in the midgut, causing similar pathologies to those seen in starvation (Christian, 1987; Arnold *et al*., 2013; Chtarbanova *et al*., 2014; Vale & Jardine, 2017). DCV shows specific tissue tropism in *D. melanogaster*, with infection of the heart tissue, fat body, visceral muscle cells around the midgut and food storage organ (crop) causing reduced defecation, food blockage and dehydration/starvation (Ferreira *et al*., 2014). Infection progresses in a similar manner following both oral or septic inoculation, with the same tissues ultimately becoming infected (Cherry & Perrimon, 2004; Arnold *et al*., 2013; Chtarbanova *et al*., 2014; Ferreira *et al*., 2014). If hosts are in a nutritional environment that allows for investment in immune function or damage repair, they may be more able to resist, or tolerate a novel infection (Ponton *et al*., 2011, 2013; Pike *et al*., 2019). This could then lead to different outcomes following a host shift, either the host could manage to suppress the parasite or avoid infection entirely, or could become infected and minimise parasite damage (Lazzaro & Little, 2009; Howick & Lazzaro, 2014). Alternatively hosts may be fully susceptible to infection, and enriched resources may act to enhance pathogen virulence by enabling within host pathogen growth (Hall *et al*., 2009; Pike *et al*., 2019). Previous work has demonstrated that following inoculation into a novel host species, the host phylogeny is an important determinant of susceptibility to DCV (Longdon *et al*., 2011, 2015). The host phylogeny explains a large proportion of the variation in DCV virulence (mortality) and viral load (75% and 67% respectively) with high virulence being associated with high viral loads (Longdon *et al*., 2015). One of the fundamental steps needed for a successful host shift is the ability of a pathogen to infect a novel host. Here we ask if the nutritional environment alters the susceptibility to DCV following a shift into a range of novel host species, and whether such patterns are consistent across species.

## Methods

### Diet preparation

Three different cornmeal diets were used (*Supplementary of species used and food type*). The standard cornmeal diet used in our lab comprised a 1:10 protein to carbohydrate ratio and became our “Medium” -protein: carbohydrate ratio diet treatment. We also developed two further diets that were approximately isocaloric, a low protein: carbohydrate food (1: 20 protein to carbohydrate) and a high protein: carbohydrate food (1: 5 protein carbohydrate); see supplementary for full table of food recipes nutrient breakdown. These were based around previous findings that suggested that in *D. melanogaster* lifespan was maximized on a protein: carbohydrate ratio of around 1:16, and fitness - measured as lifetime egg production at a ratio of 1:4 (Lee *et al*., 2008). All diets were manipulated by altering the dextrose and yeast amounts whilst maintaining as close as possible the same calorie content at 142 Calories g/100 ml. Yeast was manipulated as it provides the majority of the protein as well as other non-caloric nutritional requirements (Piper, 2017). Values were confirmed using the Drosophila Dietary Composition Calculator (Lesperance & Broderick, 2020).

### Viral Infections

Twenty-seven different species of Drosophilidae were maintained in multi generation populations, in Drosophila stock bottles (Fisherbrand) on 50 ml of their respective food medium at 22°C and 70% relative humidity with a 12-hour light-dark cycle (See *Supplementary for species and food*). Each day, two vials of 0-1 day old male flies were randomly assigned to one of three potential food types; low, medium or high, protein: carbohydrate ratio. The mating status of flies was not controlled as some species may reach sexual maturity before collection. We used male flies only for this study to remove any potential effect of sex. Flies were kept on their respective food treatments for 5 days, and tipped onto fresh vials of food every day (Broderick & Lemaitre, 2012; Blum *et al*., 2013). After 5 days of acclimatisation on their food treatment flies were experimentally infected with DCV. These collections and inoculations were carried out over three replicate blocks, with each block being completed over consecutive days (Figure 1). The order that the species were infected was randomized each day, as was food treatment for each species.

**Figure 1.**
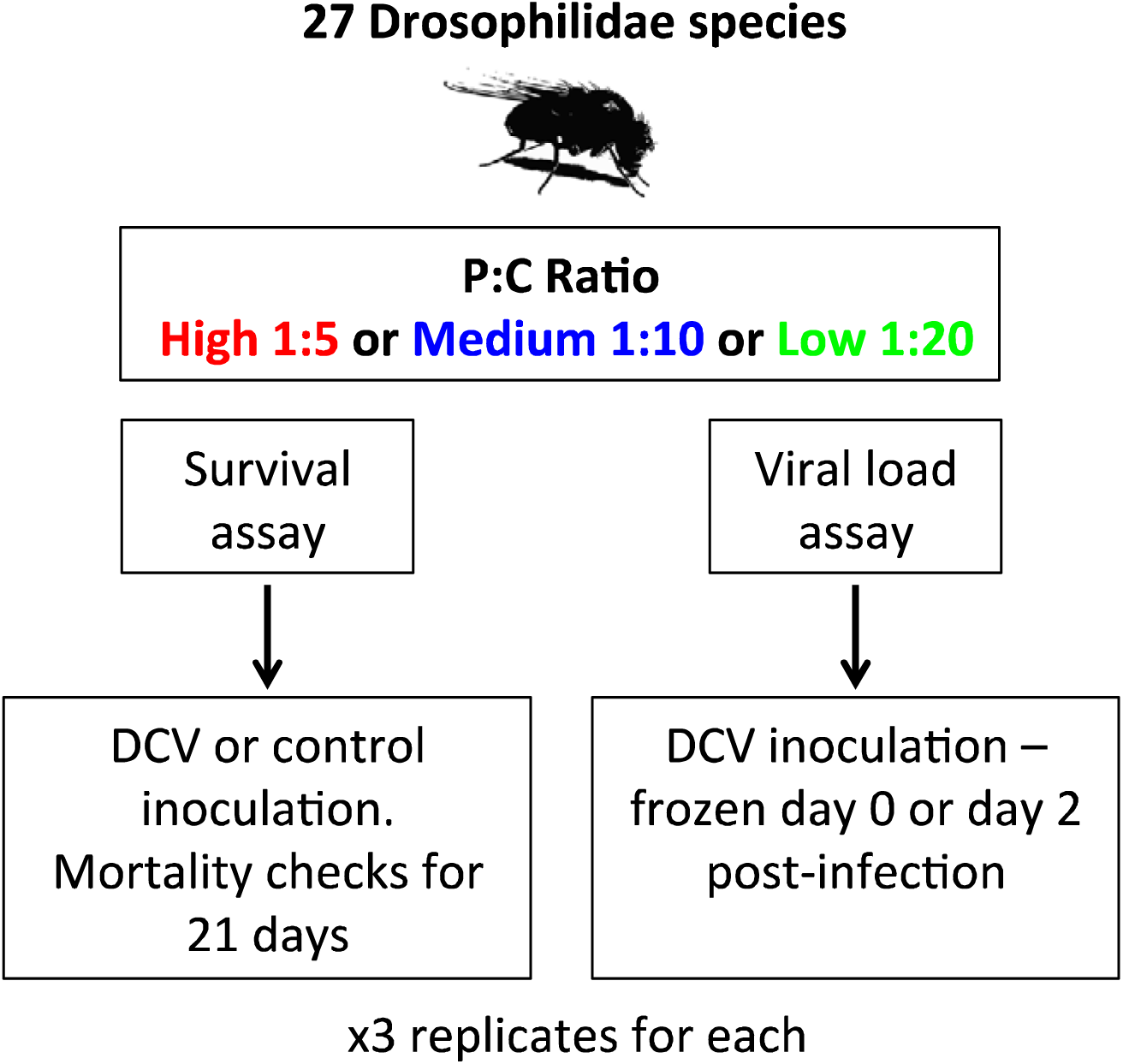
Schematic of the experimental set up: males from 27 species of Drosophilidae were housed on three foods with different protein: carbohydrate ratios before being inoculated with DCV or a control for a survival assay. Flies were also inoculated with DCV and sampled immediately (day 0) or 2 days post infection to measure the change in RNA viral load. For each treatment (control/virus or day 0/day 2) there were three replicates, where each replicate was a vial of flies (number of flies per vial described below)

We used Drosophila C virus (DCV) strain B6A (Longdon *et al*., 2018), which is derived from an isolate collected from *D. melanogaster* in Charolles, France (Jousset *et al*., 1972). The virus was prepared as described previously (Longdon *et al*., 2013). Briefly, DCV was grown in Schneider’s Drosophila line 2 cells and the Tissue Culture Infective Dose 50 (TCID50) per ml was calculated using the Reed-Muench end-point method. Flies were anesthetized on CO_2_ and inoculated using a 0.0125 mm diameter stainless steel needle that was bent to a right angle ∼0.25mm from the end (Fine Science Tools, CA, USA). The bent tip of the needle was dipped into the DCV solution (TCID50 = 6.32×10^9^) and pricked into the pleural suture on the thorax of the flies (Longdon *et al*., 2015). We selected this route of infection as oral inoculation has been shown to lead to stochastic infection outcomes in *D. melanogaster*, with injection producing a more reproducible infection, that has been found to follow a similar course to an oral infection, with the same tissues ultimately becoming infected by both methods (Cherry & Perrimon, 2004; Chtarbanova *et al*., 2014; Ferreira *et al*., 2014; Merkling & van Rij, 2015). One vial of inoculated flies was immediately snap frozen in liquid nitrogen to provide a time point zero samples to be used as a reference sample to control for relative viral dose. The second vial of flies were infected and then placed back into a new vial of their respective food treatment. After 2 days (+/- 1 hour) flies were snap frozen in liquid nitrogen. This time point was chosen based on previous studies that show a clear increase in viral growth but little mortality at 2 days post infection (Longdon *et al*., 2015; Roberts *et al*., 2018). Each experimental block contained a day 0 and day 2 replicate for each species, at each diet (27 species × 3 diet treatments × 3 experimental blocks). In total, we quantified viral load in 7580 flies in 474 biological replicates (biological replicate = change in viral load from day 0 to day 2 post-infection), with a mean of 16 flies per replicate (range across species = 8-28).

### Survival

In order to measure the effect of diet on virulence we also carried out a survival assay where mortality was recorded following infection. The same infection protocol was carried out as above; one vial of flies was infected with DCV whilst the other was injected using a clean virus free needle dipped in *Drosophila* Ringer’s solution (Sullivan *et al*., 2000) (Figure 1). Flies were maintained in vials as described above and tipped onto their respective fresh food every 2 days. The number of dead flies was counted every day for 21 days. The survival assay was carried out across three blocks with infections carried out over consecutive days, to obtain a control and infected vial per species each day. Treatment (virus or control) and the order in which fly species were inoculated were randomized between blocks. Diet was randomized across days, so for a given food type a control and viral infected vial was completed each day, and this was repeated over consequent days until there was a control and infected for each species on each food type (27 species × 2 treatments (control or challenged) × 3 diet treatments × 3 experimental blocks). In total, we measured mortality in 9222 flies with a mean of 20 flies per replicate (range across species: 6–30 flies).

### Measuring the change in viral load

The change in RNA viral load from day 0 to day 2-post infection was measured using quantitative Reverse Transcription PCR (qRT-PCR). Frozen flies were homogenised in Trizol reagent (Invitrogen) using a bead homogeniser for 30 seconds (Bead Ruptor 24; Omni international) and stored at −80°C for later extraction. Total RNA was extracted from the Trizol homogenised flies in a chloroform isopropanol extraction, reverse-transcribed with Promega GoScript reverse transcriptase and random hexamer primers. Viral RNA load was expressed relative to the endogenous control housekeeping gene *RpL32*. Primers were designed to match the homologous sequence in each species and crossed an intron-exon boundary so will only amplify mRNA. The primers in *D. melanogaster* were *RpL32* qRT-PCR F (5′-TGCTAAGCTGTCGCACAAATGG −3′) and *RpL32* qRT-PCR R (5′-TGCGCTTGTTCGATCCGTAAC −3′) (see supplementary table and Longdon *et al*., 2011). DCV primers were 599F (5′-GACACTGCCTTT GATTAG-3′) and 733R (5′CCCTCTGGGAACTAAATG-3′) as previously described (Longdon *et al*., 2015). Two qRT-PCR reactions (technical replicates) were carried out per sample with both the viral and endogenous control primers, with replicates distributed across plates in a randomised block design. qRT-PCR was performed on an Applied Biosystems StepOnePlus system using Sensifast Hi-Rox Sybr kit (Bioline) with the following PCR cycle: 95°C for 2 min followed by 40 cycles of: 95°C for 5 sec followed by 60°C for 30 sec. Each qRT-PCR plate contained four standard samples. A linear model was used to correct the cycle threshold (Ct) values for differences between qRT-PCR plates. Samples where the technical replicates had Ct values more than 2 cycles apart after plate correction were repeated. To estimate the change in viral load, we first calculated ΔCt as the difference between the cycle thresholds of the DCV qRT-PCR and the RpL32 endogenous control. For each species the viral load of day 2 flies relative to day 0 flies was calculated as 2^-ΔΔCt^; where ΔΔCt = ΔCt day0 – ΔCt day2. The ΔCt day 0 and ΔCt day 2 is a pair of ΔCt values from a day 0 biological replicate and a day 2 replicate.

**Table 1.** Ingredients for the experimental diet treatments. Amounts given are enough to produce ∼100 vials of food, with calculated calories per 100ml.

| Diet | Ratio Protein: Carb | Cornmeal (g) | Dextrose (g) | Yeast (g) | Agar (g) | Nipagin (ml) | dH2O (L) | Calories per 100ml |
| --- | --- | --- | --- | --- | --- | --- | --- | --- |
| High | 1:5 | 176 | 131.2 | 84 | 22 | 29 | 1 | 142.60 |
| Medium | 1:10 | 176 | 176 | 38 | 22 | 29 | 1 | 142.30 |
| Low | 1:20 | 176 | 203 | 10 | 22 | 29 | 1 | 142.02 |

### Effect of Body Size

To account for any potential differences in body size between species, we measured wing length as a proxy for body size (Huey *et al*., 2006). During the collections for the viral load assay males of each species were collected and immediately stored in ethanol. Subsequently, wings were removed and photographed under a dissecting microscope. Using ImageJ software (version 1.48) the length of the IV longitudinal vein from the tip of the proximal segment to where the distal segment joins vein V was recorded, and the mean taken for each species, overall there was a mean of 28 wings measured per species (range 20–35).

### Host phylogeny

The host phylogeny was inferred as described previously (Longdon *et al*., 2015) using seven genes (mitochondrial; *COI, COII*, ribosomal; *28S* and nuclear; A*dh, SOD, Amyrel, RpL32*). Publicly available sequences were downloaded from Genbank or were Sanger sequenced. In total we had *RpL32* sequences for all 27 species, *28S* from 24 species, *Adh* from 24 species, *Amyrel* from 15 species, *COI* from 27 species, *COII* from 27 species and *SOD* from 12 species. For each gene the sequences were aligned in Geneious (version 9.1.8) (Kearse *et al*., 2012) using the global alignment setting, with free end gaps and a cost matrix of 70% similarity. The phylogeny was inferred using BEAST (v1.10.4) (Drummond *et al*., 2012), with genes partitioned into three groups; mitochondria, ribosomal and nuclear, with their own molecular clock models. A random starting tree was used, with a relaxed uncorrelated lognormal molecular clock. Each of the partitions used a HKY substitution model with a gamma distribution of rate variation with 4 categories and estimated base frequencies. Additionally, the mitochondrial and nuclear data sets were partitioned into codon positions 1+2 and 3, with unlinked substitution rates and base frequencies across codon positions. The tree-shape prior was set to a birth-death process. The BEAST analysis was run twice to ensure convergence for 1000 million MCMC generations sampled every 10000 steps. On completion the MCMC process was examined using the program Tracer (version 1.7.1) (Rambaut *et al*., 2014) to ensure convergence and adequate sampling, and the constructed tree was then visualised using FigTree (v1.4.4) (Rambaut, 2006).

### Statistical analysis

We used phylogenetic mixed models to look at the effects of host relatedness on mortality and viral load across the three diet treatments. We fitted all models using a Bayesian approach in the R package MCMCglmm version 2.29 (Hadfield, 2010) in RStudio (R version 3.5.1) (R Development Core Team, 2005). We used a multivariate model with mortality of the controls, mortality of the virus infected flies and viral load at each of the diets as the response variables. Mortality was calculated as the mean portion of flies alive each day for each vial. The model took the following form:

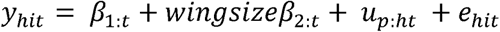

Where is the change in viral load of the *i*^th^biological replicate of host species *h*, for trait *t. β* are the fixed effects, with *β*_1_ being the intercepts for each trait and *β*_2_ being the effect of wing size. *U*_*p*_ are the random phylogenetic species effects and *e* the model residuals. The models were also run with a species-specific component independent of phylogeny (*u*_*s*:*ht*_) that allow us to estimate the proportion of variation that is not explained by the host phylogeny (*v*_*s*_) (Longdon *et al*., 2011). The main model was run without this term as it struggled to separate the phylogenetic and non-phylogenetic terms. Our main model therefore assumes a Brownian motion model of evolution (Felsenstein, 1973). The random effects and the residuals are assumed to be multivariate normal with a zero mean and a covariance structure **V**_**p**_ ⦻ **A** for the phylogenetic affects and **V**_**e**_ ⦻ **I** for the residuals (⦻ here is the Kronecker product). **A** is the phylogenetic relatedness matrix, **I** is an identity matrix and the *V* are 9_×_9 (co)variance matrices describing the (co)variances between viral load, mortality in virus infected, and mortality in controls each at the 3 different diet levels. The phylogenetic covariance matrix, **V**_**p**_ – describes the inter-specific variances in each trait and the inter-specific covariances between them. The residual covariance matrix, **V**_**e**_ describes the within-species variance that includes both any actual within-species effects and also any measurement or experimental error. The off-diagonal elements in **V**_**e**_ (the covariances) are unable to be estimated because no vial has multiple measurements – so were set to zero. The MCMC chain was run for 1300 million iterations with a burn-in of 30 million iterations and a thinning interval of 1 million. Results were tested for sensitivity to the use of different priors by being run with different prior structures (see supplementary R code), which gave qualitatively similar results. We also ran models with the data subset into viral load and mortality that gave similar results. All confidence intervals (CI’s) reported are 95% highest posterior density intervals. In order to test for the interaction between diet and species we calculated correlations between traits from the variance covariance matrix from the diet:species random effect (*u*_*p*:h*t*_). If the correlations between traits are close to one and there is no change in the means or the variance, it would suggest that there is no species-by-diet interaction. We confirmed our experimental design and sample sizes had sufficient power to detect effects by down sampling a similar dataset (Roberts *et al*., 2018).

## Results

In order to investigate the effect that host diet may have on the likelihood of virus host shifts we quantified DCV viral load in 27 species of Drosophilidae that had been housed on three different diets (Fig.2). Viral loads differed between species, with a billion times more virus in the most susceptible compared to the least susceptible species, consistent with previous studies (Longdon *et al*., 2015; Roberts *et al*., 2018). Viral loads were highly repeatable, with the inter-specific phylogenetic component (*v*_p_), explaining a high proportion of the variation in viral load across diets with little within species or measurement error (*v*_*e*_) (Repeatability = *v*_*p*_ /(*v*_*p*_ + *v*_*e*_); Low = 0.92 (95% CI: 0.86, 0.96); Medium = 0.90 (95% CI: 0.84,0.96); High = 0.83 (95% CI: 0.75, 0.92).

**Figure 2.**
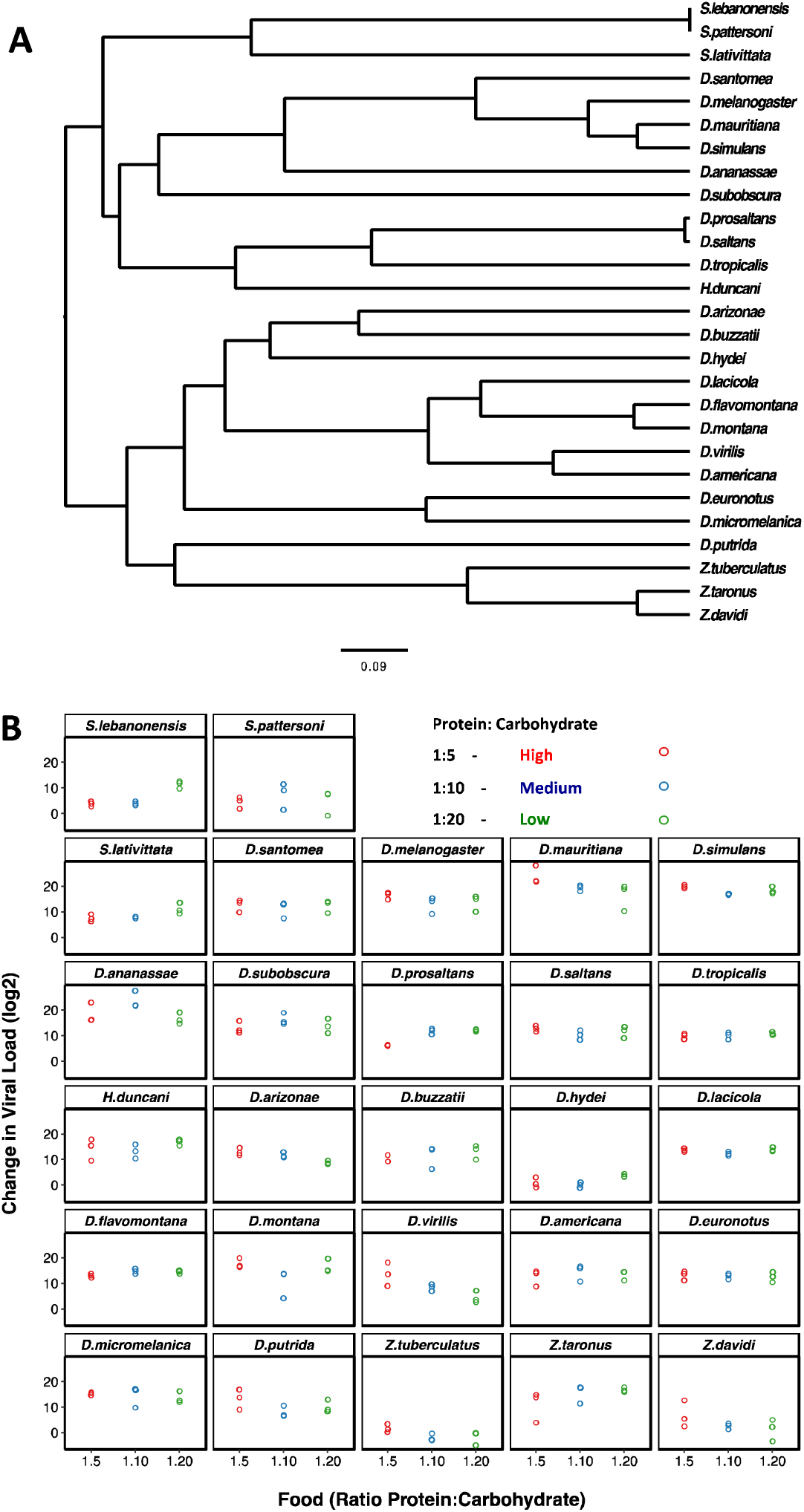

(A). Phylogeny of the 27 Drosophilidae host species. Scale bar is the number of substitutions per site. (B) Change in RNA viral load (log_2_) for the host species infected with DCV across the three different diets of differing protein: carbohydrate ratios. Individual points are plotted with a small x-axis jitter and represent the change in viral load between day 0 and day 2-post infection. Panels are ordered as on the tips of the phylogeny in (A).

We also partitioned the inter-specific variance into that which can be explained by a Brownian motion model of evolution on the host phylogeny (*v*_*p*_), and a species-specific component independent of the phylogeny (*v*_*s*_). The proportion of the between species variance that can be explained by the phylogeny can then be calculated, using *v*_p_/(*v*_p_ + *v*_s_) (Freckleton *et al*., 2002), and can be equated to the phylogenetic heritability or Pagel’s lambda (Pagel, 1999; Housworth *et al*., 2004). We found that the host phylogeny explained a modest amount of the inter-specific variation in viral loads across diets, however these estimates had broad confidence intervals (Low = 0.20 (95% CI: 3.5 ⨯10^−6^, 0.63); Medium = 0.34 (95% CI: 2.0 ⨯ 10^−6^, 0.80); High = 0.51 (95% CI: 3.2 ⨯ 10^−6^, 0.88), due to the model struggling to separate out the phylogenetic and non-phylogenetic components.

In order to examine if the susceptibility of species responded in the same or different ways to the changes in diet we examined viral load across the different protein: carbohydrate ratios. We found strong positive inter-specific correlations between viral loads across diet treatments suggesting the species are responding in similar ways to the changes in ratios (Table 2). There was a decline in the between species variance in the high diet compared to low and medium – but this was not significantly different– (*v*_p_: Low =77.13 (95% CI: 35.09, 125.50); Medium =82.33 (95% CI: 37.55, 135.26); High =45.59 (95% CI: 19.61, 76.39) and mean viral loads were similar across the diets (Low = 11.4 (95% CI: 5.3, 17.7); Medium = 10.6 (95% CI: 3.6, 16.6); High = 10.6 (95% CI: 3.6, 16.7). Residual variance did not differ significantly between treatments (Low = 6.45 (95% CI: 4.87, 8.04); Medium = 8.23 (95% CI: 6.45, 10.35); High = 8.32 (95% CI: 6.54, 10.6).

**Table 2.**
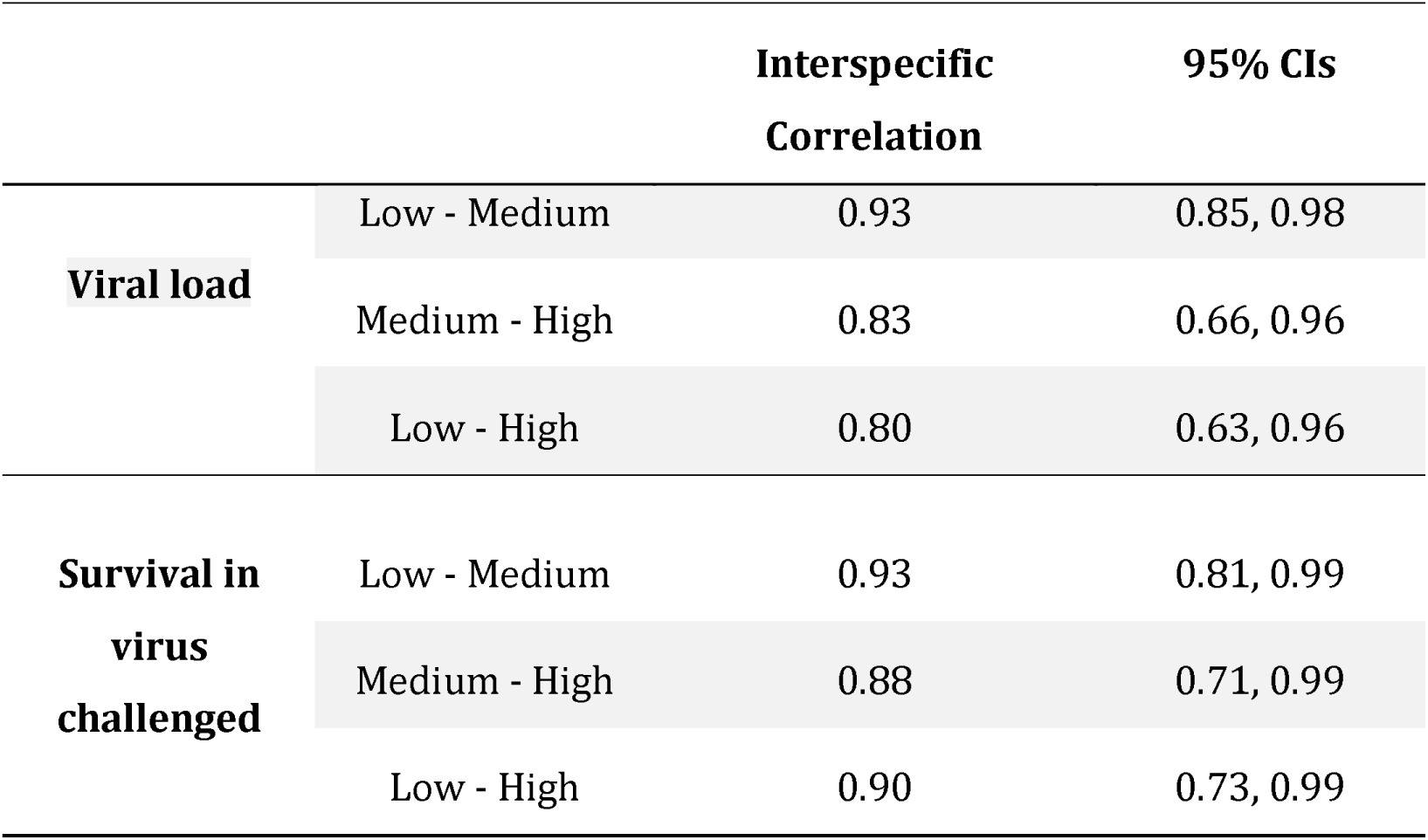
Inter-specific correlations between viral load and mortality measures across the different diet treatments.

As similar pathogen loads can cause different levels of harm to their hosts (Roy & Kirchner, 2000; Boots, 2008; Råberg *et al*., 2009) we examined if virus induced mortality differed across diets over a 20 day period after viral challenge (Fig. 3). We found differences in the virulence (mortality) caused by DCV between host species, with some species seeing no apparent change in mortality over the experimental period compared to sham infected controls, (e.g. *S. pattersoni* and *D. saltans*), whilst other species show higher susceptibility with up to 50% of flies dead by day 10 post infection (e.g. *D. simulans* and *D. melanogaster*). As with the viral load data we calculated the repeatability of survival in these virus infected flies which was high in all cases (Repeatability; Low = 0.90 (95% CI: 0.78, 0.98); Medium = 0.87 (95% CI: 0.71,0.97); High = 0.98 (95% CI: 0.85, 1.00). We also calculated the proportion of between species variance that can be explained by the phylogeny for the virus infected flies (Phylogenetic effect: Low = 0.16 (95% CI: 2.58 ⨯10^−6^, 0.62); Medium = 0.18 (95% CI: 5.75 ⨯ 10^−7^, 0.78); High = 0.32 (95% CI: 7.75 ⨯ 10^−8^, 0.87), which – like the viral load data – had broad confidence intervals due to the model struggling to separate the phylogenetic and non-phylogenetic components.

**Figure 3.**
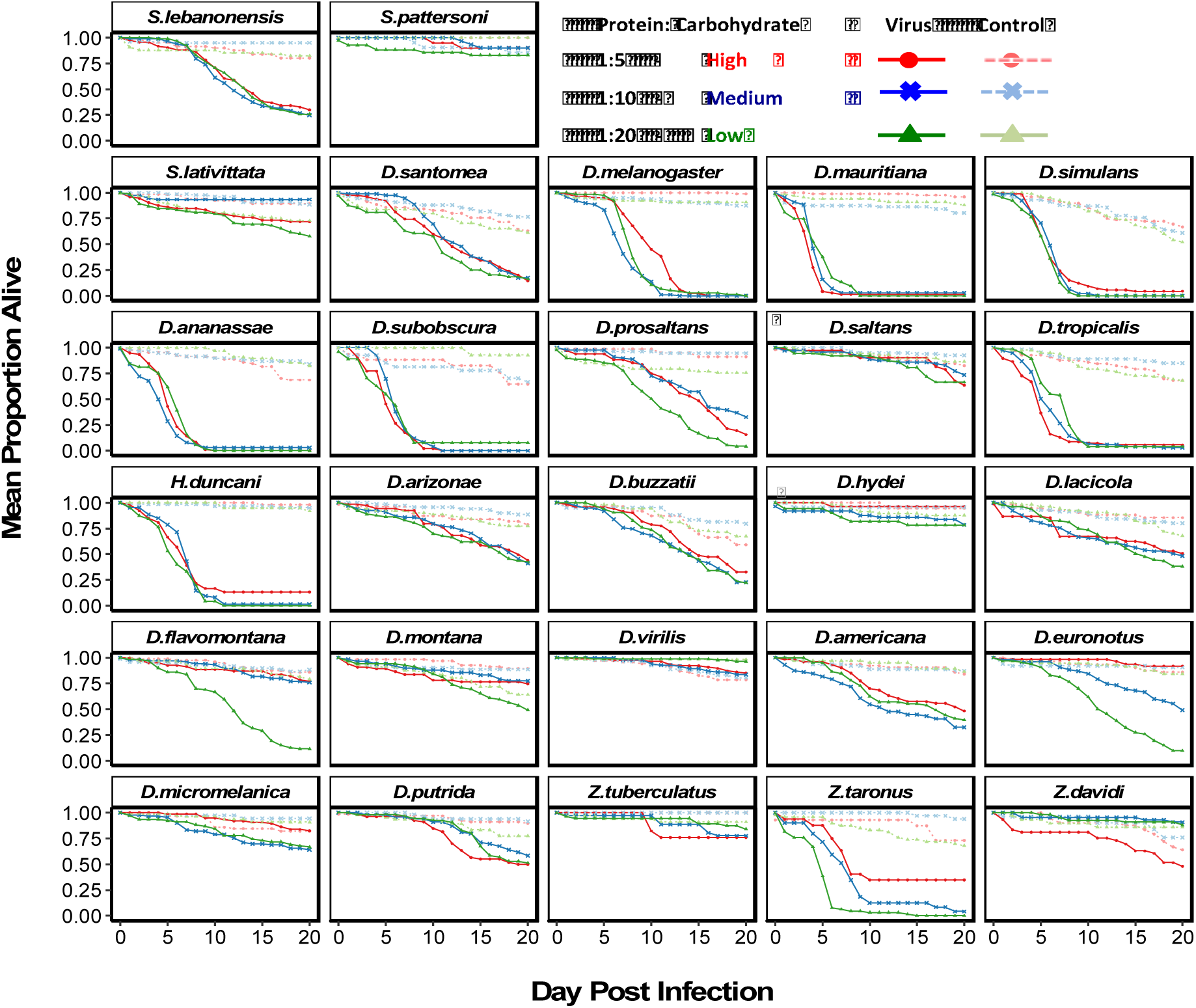
Mortality in 27 species of Drosophilidae housed on three different diets of varying protein: carbohydrate ratios. High-red circles, Medium - blue crosses and Low-green triangles and either control stabbed (dashed line) or virally challenged with DCV (solid lines). Panels are labelled in line with the tips in Figure 2A.

We found strong positive inter-specific correlations between the survival of virus challenged flies across the diets, suggesting the species are responding in similar ways to the dietary changes (Table 2). Among species variance in mortality of virus infected flies was consistent across diets (Low = 0. 18 (95% CI: 0.07, 0.31); Medium = 0.16 (95% CI: 0.04, 0.30); High = 0.12 (95% CI: 0.04, 0.23) as was the mean mortality (Low = 0.64 (95% CI: 0.47, 0.82); Medium = 0.58 (95% CI: 0.38, 0.75); High = 0.65 (95% CI: 0.47, 0.82). The residual variance was also consistent across the diets (Low = 0.02 (95% CI: 0.01, 0.03); Medium = 0.02 (95% CI: 0.01, 0.04); High = 0.02 (95% CI: 0.01, 0.03).

We found that there were strong positive correlations between mortality and RNA viral load (interspecific correlations between viral load and survival of virus infected flies: Low = 0.89 (95% CI: 0.78, 0.98); Medium = 0.85 (95% CI: 0.67, 0.97); High = 0.67 (95% CI: 0.35, 0.90). To confirm that these differences are due to mortality caused by the virus rather than intrinsic differences in the survivorship of the different species, we also inoculated flies with a control solution. There was far less mortality in the controls than the virus infected flies (Fig. 3). There was inter-specific variation in control mortality (Low = 0.18 (95% CI: 0.01, 0.59); Medium = 0.43 (95% CI: 0.01,0.76); High = 0.55 (95% CI: −0.72, 1.00) but this was not significantly correlated with survival of the virus infected flies (survival of control versus virus infected on: Low = −0.11 (95% CI: −0.92, 0.75); Medium = 0.34 (95% CI: −0.48 0.97); High = 0.12 (95% CI: −0.82, 0.89). We found no effect of wing length as a proxy for body size, (mean: −0.05, 95% CI: − 0.13, 0.05).

## Discussion

We found dietary treatments of differing protein to carbohydrate ratios did not alter the outcome of infection in 27 species of Drosophilidae infected with DCV. We found strong positive inter-specific correlations across diets in both viral load and mortality (Table 2), suggesting that the species are in general responding in similar ways to nutritional changes. Despite there being among species variation in susceptibility, generally changes in diet did not affect viral loads, nor did they alter the likelihood of surviving an infection. We found strong positive correlations between mortality and viral load on each of the diets, suggesting the amount of harm caused to a host is a result of virus accumulation within the infected host.

Although the point estimates of the inter-specific correlations are close to one (Table 2) – suggesting overall there is limited evidence for interactions between species and diet, some species do appear to show differences in mortality on different diets (e.g. *D. euronotus* and *D. flavomontana*, Figure 3). These patterns however, are not present when looking at the viral load data for these species, and our power analysis suggests we have enough power to detect interaction effects with our present experimental design. Therefore, further experiments designed to look specifically at the differences within species are required to determine if these patterns of mortality would hold true.

Both mounting and maintaining an immune response requires energy and nutrients. During an acute immune challenge the provisioning of nutrients may become more demanding for a host, with pathogen induced malabsorption through damage to or obstruction of digestive tissues (Lochmiller & Deerenberg, 2000). DCV is known to cause severe pathology of the tissues of the digestive tract with subsequent accumulation of food in the crop (food storage organ) and obstruction in the intestine (Chtarbanova *et al*., 2014). These physical symptoms alter an infected hosts energy stores with infected flies showing significantly reduced glycogen and triglyceride levels three to four days post infection (Chtarbanova *et al*., 2014). DCV infected flies also increase in body mass, with a reduced food intake and reduced metabolism, suggesting that they experience increased water retention (Thomas-Orillard, 1984; Arnold *et al*., 2013; Chtarbanova *et al*., 2014). We therefore hypothesised that changing the ratio of protein to carbohydrate in the diet may alter outcome of infection, and as species may all have their own “optimal diet”, that species may respond in different ways to such changes. However, this does not appear to be the case.

Geometric frameworks for nutrition were developed in response to the fact that what is “optimal” will depend on a balance of particular nutrients in the organism and trait being investigated (Simpson & Raubenheimer, 1995; Archer *et al*., 2009; Cotter *et al*., 2019). For example mice infected with *Salmonella* were found to survive better on diets containing a higher ratio of protein to carbohydrate (Peck *et al*., 1992). As were army worm caterpillars infected with bacteria, with survival increasing with dietary protein, suggesting high protein requirements are associated with bacterial resistance (Povey *et al*., 2009). A recent study used 10 different protein: carbohydrate diets and challenged flies with *Pseudomonas entomophila* bacteria (Savola *et al*., 2020a). Survival on low protein diets was found to be lower in infected flies and suggested protein was important for survival during infection. This study also monitored lifespan and reproduction in flies, and found that regardless of injury and infection, dietary restriction extended lifespan and reduced reproductive output (Savola *et al*., 2020a). One potential mechanism of the interaction of diet and infection has been suggested in research using a model host-pathogen system *in vivo* and *in vitro* (Wilson *et al*., 2020). Caterpillars of *S. littoralis* challenged with the bacteria *X. nematophila in vivo* and on high dietary protein had slower bacterial growth with higher survival. When this was combined with *in vitro* experiments the results suggested this was driven by the osmolality of the hosts’ blood (hemolymph) being altered by an increase in solutes in the high protein diets slowing the bacterial growth (Wilson *et al*., 2020).

Further research on the mechanistic basis of dietary effects on resistance is needed for other pathogen taxa, including viruses. Immunity to DCV inoculation in *D. melanogaster* has been reported to involve the JAK/STAT and Imd pathways, and potentially phagocytosis (Van Rij *et al*., 2006; Zhu *et al*., 2013; Lamiable *et al*., 2016). Additionally, the RNAi pathway is a key antiviral defence mechanism in *Drosophila* and DCV appears to have evolved to suppress this response (Van Rij *et al*., 2006). Although we find no interaction between dietary protein:carbohydrate and susceptibility, the multifaceted immune response to DCV may be energetically costly and other nutrients may interact with the ability of a host to allocate resources between an immune response, damage repair and the maintenance of homeostasis (Lochmiller & Deerenberg, 2000; Zuk & Stoehr, 2002; Schmid-Hempel, 2005; Sadd & Siva-Jothy, 2006). For example, lipid and fats have been associated with *D. melanogaster* response to DCV viral infection; peroxisomes were found to be required for host defense to infection, through their primary function in lipid metabolism (Aubert *et al*., 1995). The lipid level across our diets was held constant, but this may be a potential area for further study. There has been an increased use of a chemically defined (holidic) diet in order to manipulate individual nutrients present in fly diets (Lee *et al*., 2006). Exome matched diets have been shown to alleviate trade-offs in fecundity and longevity (Piper *et al*., 2017). A possible extension of this would be to look at the effect of matching diets to transcriptional changes during infection, and seeing if this alleviates (or exacerbates) pathology.

Changes in diet have been shown to alter pathogen susceptibility in a number of systems. We hypothesised that changes in diet could alter the potential outcomes of virus host shifts. However, we found that overall changes in the ratio of protein to carbohydrate did not alter susceptibility to DCV across host species in this instance. This suggests dietary protein to carbohydrate ratios are not universally important in determining susceptibility to pathogens. It is unclear if the lack of studies showing no effect of diet reflect publication biases or whether our model system is unusual. However, it highlights the need to examine the importance of diet in explaining susceptibility to pathogens across a broad range of host and pathogen taxa.

