## Supplementary Data for "Viral susceptibility across host species is largely independent of dietary protein to carbohydrate ratios"

**Supplementary Table**: List of species used in the experiment and their rearing food for stock populations. All cornmeal and Proprionic medium have dried yeast sprinkled onto the surface of the food, other food types do not unless stated below. Stock lines are the Drosophila Species Stock Centre strain IDs, Cambridge refers to flies from stocks at the Genetics Department University of Cambridge. Flies are from the following genera: Drosophila, Scaptodrosophila , Hirtodrosophila or Zaprionous.

| **Species** | **Rearing food** | **Stock line** |
| --- | --- | --- |
| *D.americana* | Malt | 15010‑0951.20 |
| *D.ananassae* | Cornmeal | Cambridge |
| *D.arizonae* | Banana | 15081‑1271.26 |
| *D.buzzatii* | Malt | 15081‑1291.01 |
| *D.euronotus* | Cornmeal | 15030‑1131.01 |
| *D.flavomontana* | Malt + yeast | 15010‑0981.05 |
| *D.hydei* | Cornmeal | 15085‑1641.71 |
| *D.lacicola* | Malt | 15010‑0991.17 |
| *D.mauritiana* | Proprionic | Cambridge |
| *D.melanogaster* | Cornmeal | Cambridge |
| *D.micromelanica* | Cornmeal | 15030‑1151.01 |
| *D.montana* | Malt + yeast | Cambridge |
| *D.prosaltans* | Proprionic | 14045‑0901.02 |
| *D.saltans* | Cornmeal | Cambridge |
| *D.santomea* | Cornmeal | 14021‑0271.00 |
| *D.simulans* | Cornmeal | Cambridge |
| *D.subobscura* | Cornmeal | 14011‑0131.15 |
| *D.virilis* | Proprionic | Cambridge |
| *H.duncani* | Proprionic | 92000‑0075.00 |
| *S. lativittata* | Banana | 11020-0081.00 |
| *S.lebanonensis* | Proprionic | Cambridge |
| *S.pattersoni* | Banana | 11010-0031.00 |
| *Z. davidi* | Banana | 50001-1040.00 |
| *Z. taronus* | Banana | 50001-1020.00 |
| *Z. tuberculatus* | Banana | 50001‑0001.05 |
| *D. putrida* | Cornmeal | 15150‑2101.00 |
| *D. tropicalis* | Cornmeal | 14030‑0801.01 |

Food recipes can be found in:

1. **Longdon B, Hadfield JD, Day JP, Smith SC, McGonigle JE, Cogni R, et al. The Causes and Consequences of Changes in Virulence following Pathogen Host Shifts. PLoS Pathog. 2015;11(3):e1004728. Epub 2015/03/17. doi: 10.1371/journal.ppat.1004728. PubMed PMID:25774803.**
2. **Longdon B, Hadfield JD, Webster CL, Obbard DJ, Jiggins FM. Host Phylogeny Determines Viral Persistence and Replication in Novel Hosts. Schneider DS, editor. PLoS Pathog. (2011) ;7: e1002260. doi:10.1371/journal.ppat.1002260**

**Supplementary Table: Food recipes and nutrient breakdown**

| Laboratory Normal Cornmeal Diet | | |  | |  |
| --- | --- | --- | --- | --- | --- |
|  | **Cornmeal** | **Dextrose (g)** | **Yeast** | **Agar** | **Totals** |
|  | **88** | **88** | **19** | **11** | **206** |
| Kcal | **325.6** | **321.2** | **67.45** | **2.86** | **717.11** |
| Fat (g) | **2.464** | **0.088** | **1.52** | **0** | **4.072** |
| Carbohydrate (g) | **69.52** | **83.6** | **3.61** | **0.77** | **157.5** |
| Protein (g) | **6.16** | **0** | **8.265** | **0.055** | **14.48** |
| Salt (g) | **0** | **0** | **0.024795** | **0** | **0.024795** |
| Sugars (g) | **1.408** | **83.6** | **2.66** | **0.033** | **87.701** |

| Diet | Ratio Protein: Carb | Cornmeal (g) | Dextrose (g) | Yeast (g) | Agar (g) | Nipagin (ml) | dH20 (L) | Calories per 100ml |
| --- | --- | --- | --- | --- | --- | --- | --- | --- |
| High | **1:06** | **176** | **131.2** | **84** | **22** | **29** | **1** | **142.60** |
| Medium | **1:10** | **176** | **176** | **38** | **22** | **29** | **1** | **142.30** |
| Low | **1:20** | **176** | **203** | **10** | **22** | **29** | **1** | **142.02** |

**Supplementary Table:** q-PCR RpL32 primer sequences for the fly species used in the experiment

| Primer Type | Primer Name | Sequence |
| --- | --- | --- |
| Forward Primers | F-a | TGCCAAGTTGTCGCACAAATGG |
|  | F-b | TGCTAAGTTGTCGCACAAATGG |
|  | F-c | TGCCAAGCTGTCGCACAAATGG |
|  | F-d | TGCTAAGCTGTCGCACAAATGG |
|  | F-e | TGCGAAGTTGTCGCACAAATGG |
|  | F-f | TGCGAAGCTGTCGCACAAATGG |
| Reverse Primers | R-a | TGCGCTTGTTGGAACCGTAAC |
|  | R-b | TGCGCTTGTTGGATCCGTAAC |
|  | R-c | TGCGCTTGTTGGAACCATAAC |
|  | R-d | TGCGCTTGTTGGAGCCGTAAC |
|  | R-e | TGCGCTTGTTAGAACCGTAAC |
|  | R-f | TACGCTTGTTGGAACCGTAAC |
|  | R-g | TGCGCTTGTTGGAACCGTAGC |
|  | R-h | TGCGCTTGTTCGATCCGTAAC |
|  | R-i | TGCGCTTGTTGGAGCCATAAC |
|  | R-j | TGCGCTTGTTTGATCCGTAAC |
|  | R-k | TGCGCTTGTTTGAACCATAAC |
|  | R-l | TACGCTTGTTGGAACCATAAC |
|  | R-m | TACGCTTGTTGGAGCCGTAAC |
|  | R-n | TGCGCTGGTTGGAACCATAAC |
|  | R-o | TGAGCTTGTTCGATCCGTAAC |
|  | R-p | TACGCTTGTTGGAGCCATAAC |
|  | R-q | TGAGCTTGTTTGATCCGTAAC |
|  | R-r | TAAGCTTGTTGGATCCGTAGC |
|  | R-s | TCAGCTTGTTGGATCCATAGC |

**Supplementary Table:** q-PCR RpL32 primer pairs for the fly species from the above sequence table

|  | ***Species*** | ***F-primer*** | ***R-primer*** |
| --- | --- | --- | --- |
| 1 | *Zaprionus davidi* | F-a | R-c |
| 2 | *Zaprionus taronus* | F-a | R-c |
| 3 | *Zaprionus tuberculatus* | F-a | R-c |
| 4 | *Drosophila putrida* | F-d | R-q |
| 5 | *Drosophila micromelanica* | F-a | R-g |
| 6 | *Drosophila euronotus* | F-a | R-g |
| 7 | *Drosophila americana* | F-c | R-a |
| 8 | *Drosophila virilis* | F-c | R-a |
| 9 | *Drosophila montana* | F-c | R-a |
| 10 | *Drosophila flavomontana* | F-c | R-a |
| 11 | *Drosophila lacicola* | F-c | R-a |
| 12 | *Drosophila hydei* | F-a | R-a |
| 13 | *Drosophila buzzatii* | F-a | R-e |
| 14 | *Drosophila arizonae* | F-a | R-a |
| 15 | *Hirtodrosophila duncani* | F-f | R-c |
| 16 | *Drosophila tropicalis* | F-b | R-k |
| 17 | *Drosophila saltans* | F-a | R-n |
| 18 | *Drosophila prosaltans* | F-a | R-n |
| 19 | *Drosophila subobscura* | F-a | R-i |
| 20 | *Drosophila ananassae* | F-f | R-a |
| 21 | *Drosophila simulans* | F-d | R-h |
| 22 | *Drosophila mauritiana* | F-d | R-h |
| 23 | *Drosophila melanogaster* | F-d | R-h |
| 24 | *Drosophila santomea* | F-a | R-n |
| 25 | *Scaptodrosophila lativittata* | F-a | R-m |
| 26 | *Scaptodrosophila lebanonensis* | F-d | R-h |
| 27 | *Scaptodrosophila pattersoni* | F-a | R-m |

**Supplementary Figure: Phylogeny of the 27 host species as constructed by BEAST program,** node labels on the phylogeny are posterior supports and the scale bar is the number of substitutions per site

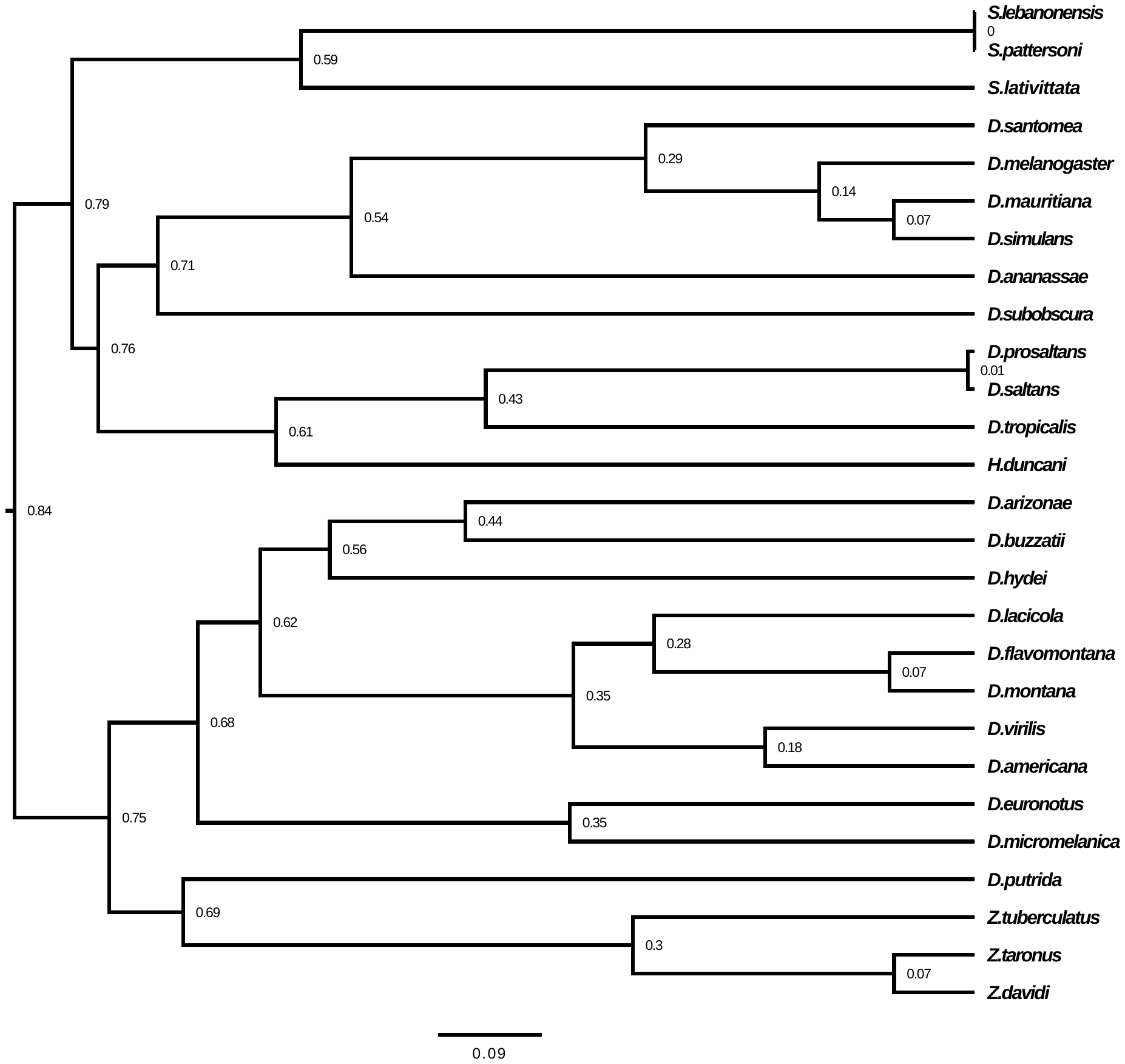

**Full project dataset: figshare.com/**

- Genbank accession numbers of sequences used to infer the host phylogeny: 10.6084/m9.figshare.13079366
- Full dataset for main analysis:
- R code: https://doi.org/10.6084/m9.figshare.13079351.v1
- Phylogenetic Tree for analysis: 10.6084/m9.figshare.13079402
